# Bridging sensory and language theories of dyslexia: towards a multifactorial model

**DOI:** 10.1101/773853

**Authors:** Gabrielle O’Brien, Jason Yeatman

## Abstract

Competing theories of dyslexia posit that reading disability arises from impaired sensory, phonological, or statistical learning mechanisms. Importantly, many theories posit that dyslexia reflects a cascade of impairments emanating from a “core deficit”. Here we collect a battery of psychophysical and language measures in 106 school-aged children to investigate whether dyslexia is best conceptualized under a core-deficit model, or as a disorder with heterogenous origins. Specifically, by capitalizing on the drift diffusion model to separate sensory encoding from task-related influences on performance in a visual motion discrimination experiment, we show that deficits in motion perception, decision making and phonological processing manifest largely independently. Based on statistical models of how variance in reading skill is parceled across measures of sensory encoding, phonological processing and decision-making, our results challenge the notion that a unifying deficit characterizes dyslexia. Instead, these findings indicate a model where reading skill is explained by several distinct, additive predictors, or risk factors, of reading (dis)ability.

**Research Highlights:**

- Our research provides direct evidence that a single-mechanism, or core-deficit, model of dyslexia cannot account for the range of linguistic and sensory outcomes in children.
- Individual differences in visual motion processing, perceptual decision making, phonological awareness and rapid naming each account for unique variance in reading skill.
- Our data support an additive risk-factor model, in which multiple independent dimensions each confer risk for reading difficulties.

## Background

Recently, there has been growing adoption of the view that dyslexia, a reading disability, is probabilistic in nature: children with a family history of dyslexia are considered “at-risk”, and compensatory skills such as strong oral language or executive functions may be “protective factors” (Haft, Myers, & Hoeft, 2016; Hulme, Nash, Gooch, Lervåg, & Snowling, 2015; Muter & Snowling, 2009; Pennington, 2006). In this multifactorial framework, most cases of dyslexia cannot be explained by a single cognitive deficit. Despite this heterogeneity, it is broadly accepted that phonological awareness (PA) and rapid automatized naming (RAN) are two of the strongest—if imperfect—predictors of reading development (Pennington et al., 2012; Wolf & Bowers, 2000).

In parallel, there is a broad literature characterizing dyslexia as the consequence of a fundamental deficit that supersedes phonological processing. There are many reports indicating that people with dyslexia also perform worse in experiments targeting various aspects of visual (Stuart, McAnally, McKay, Johnston, & Castles, 2006; Talcott et al., 2002) and auditory processing (Hämäläinen, Salminen, & Leppänen, 2013; Noordenbos & Serniclaes, 2015), as well domain general mechanisms such as processing speed and statistical learning (Gabay, Thiessen, & Holt, 2015; Vandermosten, Wouters, Ghesquière, & Golestani, 2018). These findings have spurred competing theories that explain dyslexia as the consequence of cascading effects from a fundamental sensory processing deficit. Generally, these “cascading deficit” theories contend that relatively low- or mid-level aspects of sensory processing disrupt the development of phonological awareness, and by this mechanism disrupt reading development.

Notably, these two branches of research remain largely distinct: while multifactorial models of reading disability are increasingly accepted among researchers studying high-level cognitive and linguistic functions, these models largely ignore lower-level deficits in sensory processing. In the sensory-processing literature, on the other hand, cascading deficit models continue to dominate and appeals to a “core mechanism” of dyslexia are still commonplace. Indeed, a PubMed search for the phrase “core deficit of dyslexia” turns up 118 results from 1986 to the present. Presently, hypotheses positing a core deficit with cascading effects are the focus of many neuroscientific and psychophysical studies of reading disability (Casini, Pech-Georgel, & Ziegler, 2018; Colling, Noble, & Goswami, 2017; Frey, François, Chobert, Besson, & Ziegler, 2019; Frey, François, Chobert, Velay, et al., 2019; Gori, Seitz, Ronconi, Franceschini, & Facoetti, 2016; Krause, 2015; Lieder et al., 2019; Nicolson & Fawcett, 2018; Vidyasagar, 2019).

A core deficit model is inherently at odds with a multifactorial model; to accept both models implies that a deficit is not really “core” in the majority of individuals with dyslexia. Reconciling the many disparate theories of reading disability remains a formidable challenge. To further compound the difficulty, there are several variants of the cascading deficit theory: one is the magnocellular deficit theory of dyslexia, in which a low-level impairment in the motion-sensitive magnocellular pathway of the visual system is said to disrupt reading skill development (Stein, 2001, 2018; Stein & Walsh, 1997). Proponents of this theory have argued that sensitivity to transient sensory information may not be restricted to vision, but could also affect auditory processing (Stein & Talcott, 1999; Van Ingelghem et al., 2001; Witton et al., 1998). Hypothetically, insensitivity to rapid auditory cues could diminish an individual’s ability to learn the sounds of their language (phonemes), and hence develop PA. A recent spin on this theory is the temporal processing hypothesis, which contends that, in fact, *slow* temporal mechanisms involved in entraining to the envelope of speech are the fundamental disorder (Casini et al., 2018; Goswami, 2015; Huss, Verney, Fosker, Mead, & Goswami, 2011). Distinct from these sensory theories, proponents of the statistical-learning hypothesis argue that a domain-general deficit in sensory learning and perceptual decision-making could explain why people with dyslexia perform poorly on myriad psychophysical tasks (Ahissar, 2007; Nicolson & Fawcett, 2018; Ziegler, 2008). It also purports to explain why children with dyslexia struggle to learn the mapping between letters and sounds.

Today, the literature remains inconclusive for several reasons. First, the various cascading deficit models contradict one another as each posits distinct mechanisms for disrupting phonological processing and, in turn, reading. While a statistical learning model of dyslexia could potentially explain why so many struggling readers also perform poorly on visual psychophysics, it has not been established whether these two types of deficits occur in the same individuals. The widespread use of group-level statistics makes it challenging to interpret how many individuals show a given pattern of low-level deficits, and the few studies focusing on individual patterns across a battery of diverse tasks do not encourage much hope for a uniform profile (Amitay, Ben-Yehudah, Banai, & Ahissar, 2002; Ho, Chan, Tsang, & Lee, 2002; Menghini, Carlesimo, Marotta, Finzi, & Vicari, 2010; Ramus et al., 2003; White et al., 2006).

Perhaps more importantly, it remains challenging to understand what relationship predictors from psychophysical tasks have with phonological predictors in determining reading ability--in other words, whether the influence of low-level sensory processing mechanisms on reading skill is mediated by phonological processing. Perhaps Talcott et al. (2000) best addressed this question by administering auditory, visual, and phonological tasks to 32 children, concluding that a measure of visual motion processing explained some additional variance in reading skill beyond a measure of PA. A follow-up study in more than 300 school-aged children replicated the finding that visual and auditory psychophysics explained variance in both phonological and literacy skills but did not clarify the fit of a cascading model (Talcott et al., 2002). Several others have observed evidence that psychophysical measures influence reading skill separate from the proposed phonological pathway (Snowling, Lervåg, Nash, & Hulme, 2019; Stein, 2001; White et al., 2006). Despite these findings, cascading deficit models remain at the forefront of the dyslexia debate, particularly for theories that hold a central role for sensory deficits (reviewed in (Goswami, 2015)).

There are several reasons why studies such as Talcott et al.’s are well-cited, but not broadly adopted as conclusive evidence about sensory processing in dyslexia. In the last two decades, there has been growing focus on non-sensory mechanisms that may affect how struggling readers perform on psychophysical tasks— a confound that many studies may not have sufficiently accounted for (Banai & Ahissar, 2004, 2006; Ramus & Ahissar, 2012). Furthermore, in the multifactorial literature, it is increasingly accepted that at least two dissociable aspects of phonological processing (PA and RAN) contribute to reading skill (Pennington et al., 2012; Wolf & Bowers, 1999, 2000). Most sensory literature explores the relationship of sensory measures to a single dimension of PA. As evidence mounts that PA alone is unlikely to explain many (or even most (Pennington et al., 2012)) cases of dyslexia, it remains worth considering how individual differences in visual motion processing, or perceptual decision making more generally, will fit into changing conceptions of reading disability.

Emanating from the rift in the literature, and the incompatibility of the myriad of “core deficit” models, this study investigates whether a cascading-deficit model, in which an underlying deficit in some other lower-level sensory or cognitive process disrupts phonological processing, is compatible with the pattern of behavioral testing and psychophysical results seen in a large sample of children with dyslexia. In order to separate the contributions of sensory encoding of visual motion from non-sensory aspects of the decision-making process, we revisit a widely used measure of visual motion sensitivity (random dot motion discrimination) with a mathematical modeling approach. The drift diffusion model (DDM) estimates the generating function that corresponds to an individual’s pattern of responses and reaction times on a task (Ratcliff & McKoon, 2008), and has been previously used to understand how cognitive mechanisms associated with aging (Ratcliff, Thapar, & McKoon, 2004), ADHD (Huang-Pollock et al., 2017), and development (Ratcliff, Love, Thompson, & Opfer, 2012) manifest in psychophysical task performance. The model has been extensively used to describe decision-making on the motion discrimination task (Gold & Shadlen, 2007; Palmer, Huk, & Shadlen, 2005; Shadlen, Hanks, Churchland, Kiani, & Yang, 2013), and many of its assumptions are validated by electrophysiological work in non-human primates (Shadlen & Newsome, 2001). As such, the DDM provides a rigorous way to explore the intersection of sensory integration and decision making in relation to reading skill.

Contrary to predictions of the myriad of core-deficit models, our data reveal a heterogeneity of deficits among children with dyslexia; no single factor, including measures of phonological processing, can reliably distinguish children with dyslexia from control subjects. The DDM reveals that sensory encoding and perceptual decision making are separable factors which predict unique variance in reading skill above and beyond phonological processing, and that there is not a consistent pattern of impairments among children with dyslexia. Furthermore, these sensory predictors are useful in addition to PA and RAN in characterizing an individual’s level of reading (dis)abilities. As a whole, these data provide further evidence against core-deficit models of dyslexia and indicate that multiple-deficit models must consider the combined influence of sensory, cognitive and linguistic factors on the development of reading skills.

## Methods

### Participants

A total of 119 native English-speaking school-aged children ages 8-12 were recruited for the study. Children without histories of neurological or sensory disorders were recruited from a database of volunteers in the Seattle area (University of Washington Reading & Dyslexia Research Database; http://ReadingAndDyslexia.com). Parents and/or legal guardians of all participants provided written informed consent under a protocol that was approved by the University of Washington Institutional Review Board. All subjects demonstrated normal or corrected-to-normal vision. Participants were tested on a battery of cognitive and literacy assessments, including the Woodcock-Johnson IV (WJ-IV) Letter Word Identification and Word Attack sub-tests, the Test of Word Reading Efficiency (TOWRE-2), Comprehensive Test of Phonological Processing (CTOPP-2) and the Wechsler Abbreviated Scale of Intelligence (WASI-II). All subjects had normal or corrected-to-normal vision.

Five subjects did not complete the psychophysics. An additional two subjects did not show evidence of performing above chance (greater than 60.5% accuracy at any of the four stimulus coherence levels) and were excluded from analysis. A further six subjects did not produce enough usable data to fit the DDM (no more than 15% responses outside of the acceptable response time window from 200 ms to 10 s). This left 106 subjects with usable data.

### Demographics

We recruited participants whose reading abilities ranged from profoundly impaired to highly proficient. Since reading abilities fall on a continuum (Shaywitz, Escobar, Shaywitz, Fletcher, & Makuch, 1992) we treat reading ability as a continuous measure in our main statistical analyses. For the purpose of comparison with other studies we include group-level analyses (Dyslexic versus Control) in our Supplementary Materials. Group labels were assigned on the basis of the composite Woodcock-Johnson Basic Reading Score (WJ-BRS) and TOWRE Index. As both the WJ-BRS and TOWRE Index are scored on the same standardized scale, a composite reading skill measure was created by averaging the two scores for each participant. The “Dyslexic” group comprised participants whose reading score fell 1 standard deviation or more below the population mean (reading score < 85); the “Control” group had reading skill measures above this cutoff and had never been diagnosed with a reading disability. There were 43 subjects in the Dyslexic group and 48 in the Control group. A remaining 15 subjects were not well-described by either label (e.g., reading score > 85 but an indication of a dyslexia diagnosis) so were not included in the group comparisons. As in several other studies (O’Brien, McCloy, Kubota, & Yeatman, 2018; Pennington et al., 2012), we did not IQ-match these groups, but rather controlled for nonverbal-IQ explicitly in our statistical analyses. Additionally, ADHD diagnosis was not grounds for study exclusion because of the high comorbidity between ADHD and dyslexia. The presence of ADHD was entered into our linear modeling analyses as a covariate. Relationships between demographic characteristics, phonological, IQ measures and reading skill are presented in Supplementary Tables S1 and S2.

### Healthy Brain Network dataset

The Healthy Brain Network dataset is provided to the public by the Child Mind Institute. At the time of writing, the released dataset included 1814 subjects. From this dataset, we identified 124 school-aged individuals (ages 5-17) in the urban New York City region who had been diagnosed with “Specific Learning Disorder with Impairment in Reading” by a panel of clinicians affiliated with the Child Mind Institute and had also been administered the CTOPP-2. We also identified 119 individuals who were similarly assessed and given no diagnosis of any kind. Due to the large number of participants available, we were able to create nonverbal-IQ matched control groups on the basis of the Wechsler Intelligence Scale for Children’s Matrix Reasoning scaled score (Dyslexia: *n* = 110; Control: *n* = 105). These groups did not significantly differ in terms of nonverbal-IQ (*t*(208.85) = −1.0668, *p =* 0.287) and age (*t*(212.65 = 1.041, *p =* 0.299). The Healthy Brain Network dataset can be accessed here: http://fcon_1000.projects.nitrc.org/indi/cmi_healthy_brain_network/index.html

### Psychophysics stimuli and apparatus

Stimuli for the motion discrimination experiment were created using MATLAB (The Mathworks Corporation, Natick, MA, USA) in conjunction with the Psychophysics Toolbox. Stimuli were displayed on a LG liquid crystal display (1,920 × 1,080 resolution, 120 Hz refresh rate, subtending 51° horizontally). The subjects’ response was collected using keypresses. The viewing distance was 56 cm. We used random-dot motion stimuli (150 dots) that were displayed in a circular aperture (14° in diameter) centered around the fixation mark (1°) at the center of display. Light (271 cd/m^2^) and dark (0 cd/m^2^) dots (dot size = 0.15°) moved at the speed of 8°/s on a gray background (135 cd/m^2^). Each dot was assigned a random lifetime from a uniform distribution between 0 and 200 ms (24 video frames). When a dots lifetime expired, it was randomly re-positioned within the aperture and assigned the maximum lifetime (200 ms). Motion coherence was defined as the percentage of dots moving together in the same direction compared to dots moving in random directions. The stimuli were equivalent to those used in Joo *et al*.(Joo, Donnelly, & Yeatman, 2017) except (a) with fixed coherence levels, and (b) stimuli remained on the screen until the subject indicated a decision with a button press (as opposed to fixed duration).

### Psychophysics procedure

Each session comprised 6 experimental blocks. For each subject, three blocks of fifty stimuli were tested with a brief break in between. This was followed by a longer break to collect reading, phonological and IQ measures, and followed by the final set of three blocks. At the beginning of the session, subjects completed 10 practice trials comprising high coherence motion (60–100%). Subjects were allowed to repeat the practice up to three times, until they got at least 70% correct. All participants were able to do this.

Stimuli were presented at five coherence levels: 6%, 12%, 24%, 48%, and 100%. However, early in the study we realized that many subjects (unrelated to reading ability) found 100% coherence difficult and reported varying visual percepts. Performance typically declined for 100% coherence stimuli compared to 48% coherence. Therefore, we analyzed only the range of stimulus coherence levels where performance was generally monotonic, from 6% to 48%. Each stimulus coherence level was presented 60 times for a total of 300 presentations, 240 of which were included in the analysis.

Each trial started with a fixation mark at the center of the display. After 500 ms, random-dot motion stimuli were displayed until the subject made a keypress (or until 10 seconds had elapsed). Subjects pressed right or left arrow keys on a standard keyboard to report motion direction. The fixation mark was turned off when the response was made, and visual and auditory feedback was given to indicate correct and incorrect responses. The experiment did not proceed until subjects reported the motion direction. The inter-trial interval was 1 s, and after this interval the fixation mark re-appeared at the center of the display to indicate the beginning of the next trial.

### Drift diffusion model

Fundamentally, the DDM tries to maximize the likelihood of observing a distribution of reaction times according to the probability density function

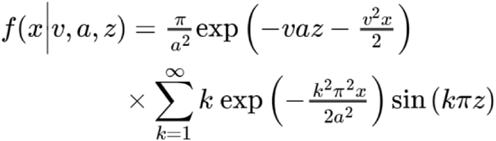

where *x* is reaction time, *v* is drift rate (the average rate at which evidence is accumulated for a decision), *a* is the distance between the two decision boundaries, and *z* is a bias term that allows for an observer to prefer one alternative to the other (Wald, 1947). The parameter *v* is allowed to vary with stimulus level. Additionally, the DDM fits a parameter *t*, which corresponds to the non-decision time- in other words, the time taken by all sensory and motor processes besides accumulating evidence for a decision, such as planning and executing a motor response and converting incoming sensory inputs to units of evidence.

The above equation predicts perfectly symmetric distributions of correct and error response times, and so the DDM model has been extended to include three free parameters that allow it to approximate more realistic distributions. The first of these parameters is *sz*, the trial-to-trial variability in the drift process starting point (centered around the halfway-point between the two decision bounds), which allows the model to predict fast errors. The other parameters are *sv*, the trial-to-trial variability in average drift rate, and *st*, the trial-to-trial variability in residual time *t*. When the DDM is fit with these three additional parameters, it is referred to as the *full* DDM.

The full DDM was fit using the Hierarchical Drift Diffusion Model toolkit for Python(Wiecki, Sofer, & Frank, 2013). Parameter reliability estimates are provided in the supplement (Table S3). The DDM was fit to each individual’s behavioral responses and reaction times using the Maximum Likelihood fitting method (as recommended by Van Zandt(2011)). The optimization scheme was attempted five times per individual and parameter estimates from the best run were saved. In all cases, the optimization scheme terminated successfully. As recommended by the makers of the HDDM package (Wiecki et al., 2013), the DDM was fit with a mixture model that allowed up to 5% of responses to be assigned to a uniform “lapse” distribution. This reduces bias in drift rate estimates due to occasional lapses. Because this mixture component was included, we employed only a coarse screen for outlier detection before DDM fitting: responses occurring before 200 ms (before a typical behavioral response can be executed) and after 10s (after the stimulus had concluded) were excluded. Although we excluded two participants with >15% data loss, the average participant in the remaining sample had 98.1% usable data.

### Outlier detection

To determine the presence of highly unusual model fits, we computed the Mahalanobis distance for each individual with respect to the 9 parameters estimated by the DDM. The Mahalanobis distance for multiple dimensions follows a chi-squared distribution, and so we use this measure to detect outliers(Filzmoser, 2004). Specifically, individuals with a Mahalanobis distance corresponding to values beyond the p < 0.001 threshold were deemed to be outliers. Two such individuals were detected; both had been fit with extremely high *a* values (*a* = 8.17 and *a* = 5.60). One of these individuals had a composite reading score in the Dyslexic range, whereas the other would have been above our cutoff. These two points were excluded from further analysis as we have cause to doubt the quality of their DDM parameter estimates, but their results are included with the full dataset online.

### Stepwise model selection procedure

In our analyses of the relationships between various parameter estimates from the DDM and reading skill, we employed a stepwise model selection procedure. In all cases, we considered three covariates: age, nonverbal-IQ, and the presence of an ADHD diagnosis. Each model selection procedure began with a fully specified model of reading score as a function of the parameter(s) of interest plus the three covariates. Fitting was performed with the base R lm() function, except where mixed model usage is noted, in which case the lme4 library was used(Bates, Sarkar, & Matrix, 2007). The contributions of the covariates were first tested using an anova test. Model terms were retained if the p-value associated with the more complex model was less than 0.1. Next, parameters of interest were tested similarly. Throughout the manuscript, wherever model selection is performed we report the selected (“most parsimonious”) model.

### Mediation analysis

Mediation analysis was performed using the “mediation” package for R(Tingley, Yamamoto, Hirose, Keele, & Imai, 2015). In all mediation models, nonverbal IQ was entered as a covariate. 4000 bootstrap simulations were used to estimate the proportion of mediation of a variable of interest by PA in a linear model of reading skill.

## Results

### Predicting dyslexia from phonological measures

We first assessed the phonological core deficit model by quantifying the extent to which deficits in PA, RAN, or both differentiate individuals with dyslexia from control subjects with typical reading skills (Figure 1). A recently released public dataset, the Child Mind Institute’s Healthy Brain Network (Alexander et al., 2017) (HBN), allows us to explore this question in a large sample of children (N = 1814, n = 110 children with dyslexia, n = 105 children selected to match on age and nonverbal IQ with no neurological or psychiatric diagnosis). Using quadratic discriminant analysis (QDA) with age and nonverbal-IQ matched groups, we asked what proportion of children could be correctly classified on the basis of two predictors: the Comprehensive Test of Phonological Processing (CTOPP-2)’s Elision measure of PA and Rapid Symbol Naming Composite measure of RAN (both age-normed). A QDA classifier trained with leave-one-out cross validation could correctly classify 67.4% (±6.3%; 95% confidence interval) of individuals with a specificity of 68.2% and a sensitivity of 66.7%. A support vector machine achieved equivalent accuracy.

**Figure 1.**
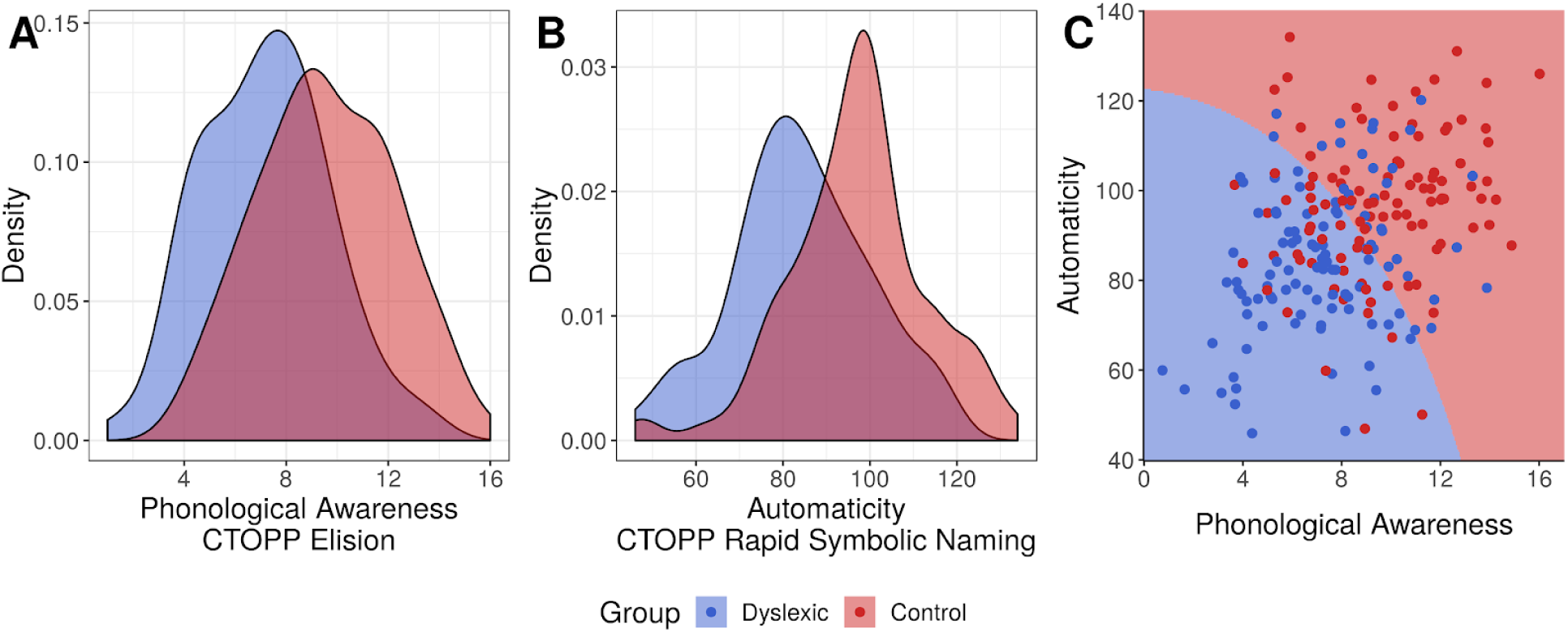
Panels A-B: Density plots for phonological awareness (PA, CTOPP Elision) and rapid automatized naming (RAN, CTOPP Rapid Symbolic Naming Composite) in the Healthy Brain Network (HBN) dataset in two groups. The Dyslexia group (blue) consists of 110 school-aged children diagnosed with dyslexia by a panel of clinicians. The red density plot represents an age- and nonverbal-IQ-matched control group of 105 children identified as having no psychiatric or neurological diagnoses by the same panel. Panel C: The decision boundary of a quadratic discriminant analysis trained on the entire dataset is shown. Dots represent observations from the dataset with slight jitter added for visibility of overlapping points.

This result is undoubtedly in alignment with the extensive literature on phonological processing: PA and RAN are both meaningful predictors of reading skill. Yet, these two measures alone fail to account for many cases of dyslexia. Furthermore, many individuals with apparently typical reading abilities would be predicted to be dyslexic on the basis of their PA and RAN scores alone.

In the original formulation of the phonological core deficit model (e.g. (Stanovich, 1988)), PA is purported to be a more powerful predictor of reading disability in early childhood, so it may be unsurprising that our model performs poorly in a sample containing teenagers. We therefore repeated our analysis on two subsets of the sample: 62 children between aged 5-8 (n = 29 with a Dyslexia diagnosis), and 153 children aged 8-17 (n = 81 with a Dyslexia diagnosis). The classifier trained on the younger cohort obtained an accuracy of 69.4% (± 11.7%) while the classifier trained on the older cohort reached 66.7% (± 7.5%). We ran a second analysis treating age as a continuous predictor: we used logistic regression on our entire sample to model dyslexia diagnosis (present or absent) with main effects of age, PA, and the interaction of the two. The interaction term was not significant (*β* = −0.007, SE = 0.0243, p = 0.767). For completeness, we tested a direct measure of pseudoword reading skill provided in the HBN dataset (the age-normed Weschler Individual Achievement Test Pseudoword subtest) as the dependent variable in a linear model. The interaction of age and PA was again not significant (*β* = −0.0322, SE = 0.116, p = 0.782). As such, our finding that standard phonological measures are only modest predictors of dyslexia in the HBN dataset is unlikely to be an artifact of the age range in the sample.

### Differences in visual motion processing

Having demonstrated that phonological predictors alone are an insufficient to explain many cases of dyslexia, we next consider the contribution of visual motion processing to reading abilities: do visual motion processing difficulties typically coincide with phonological impairments, as would be expected in a cascading model of reading disability? Or are they a separable contributor to reading outcomes which explain cases of dyslexia that were not captured by the phonological core deficit model? Here we present the results of the motion discrimination experiment (conducted in the lab) in 106 school-aged children, including 42 individuals who meet our criteria for dyslexia. Accuracy and response times were collected for stimuli presented at four coherence levels: 6%, 12%, 24%, and 48%.

Before we model the respective contributions of sensory and decision processes to task performance, it is important to establish that task performance is related to reading skill. We confirmed that reading skill was related to reaction time: using model selection, we identified that the most parsimonious model of median reaction time included main effects of stimulus coherence (*β* = −0.173, SE = 0.00898, p < 1 × 10^−15^), age (*β* = −0.059, SE = 0.0214, p < 1 × 10^−15^), and reading skill (*β* = −0.006, SE = 0.00149, *p =* 1.15 × 10^−4^) with a random effect of subject (Table S4 and Figure S1). Accuracy was not significantly related to reading skill (Table S5), likely reflecting the fact that the motion stimuli remained on the screen until the subject provided a response. Notably, we also observed that the ratio of correct to error median response times within each subject was significantly associated with reading skill (*β* = −0.00444, SE = 0.00224, *p =* 0.0497), with poor readers showing an increased tendency to make “fast errors” relative to correct response times (Table S6 and Figure S2). The presence of fast errors is notable because this phenomenon is typically associated with non-sensory mechanisms, including a tendency to initiate guesses before an optimal amount of evidence is considered (Smith & Ratcliff, 2004). Thus, raw reaction time data indicated that children with dyslexia were not only less efficient than control subjects, but also showed a qualitatively different pattern of responses.

### Less efficient visual motion processing in dyslexia

To decouple sensory encoding of visual motion from the process of forming and executing a binary decision, we fit the drift diffusion model (DDM) to each subject’s distribution of behavioral responses and reaction times. In the DDM for a two-alternative forced-choice judgment, it is assumed that an observer samples sensory input at discrete moments in time, and that these samples are accumulated in a noisy decision variable that represents the integrated evidence over the course of the trial (plus internal noise). When this decision variable reaches a threshold, the observer initiates a decision (Figure 2A). The DDM therefore separates the encoding and evaluation of sensory information (which drives changes in the decision variable) from non-sensory processes, such as the magnitude of the threshold for triggering a decision and the trial-to-trial variability in the decision process (for a detailed review of the DDM, see (Ratcliff & McKoon, 2008; Wiecki et al., 2013)).

**Figure 2.**
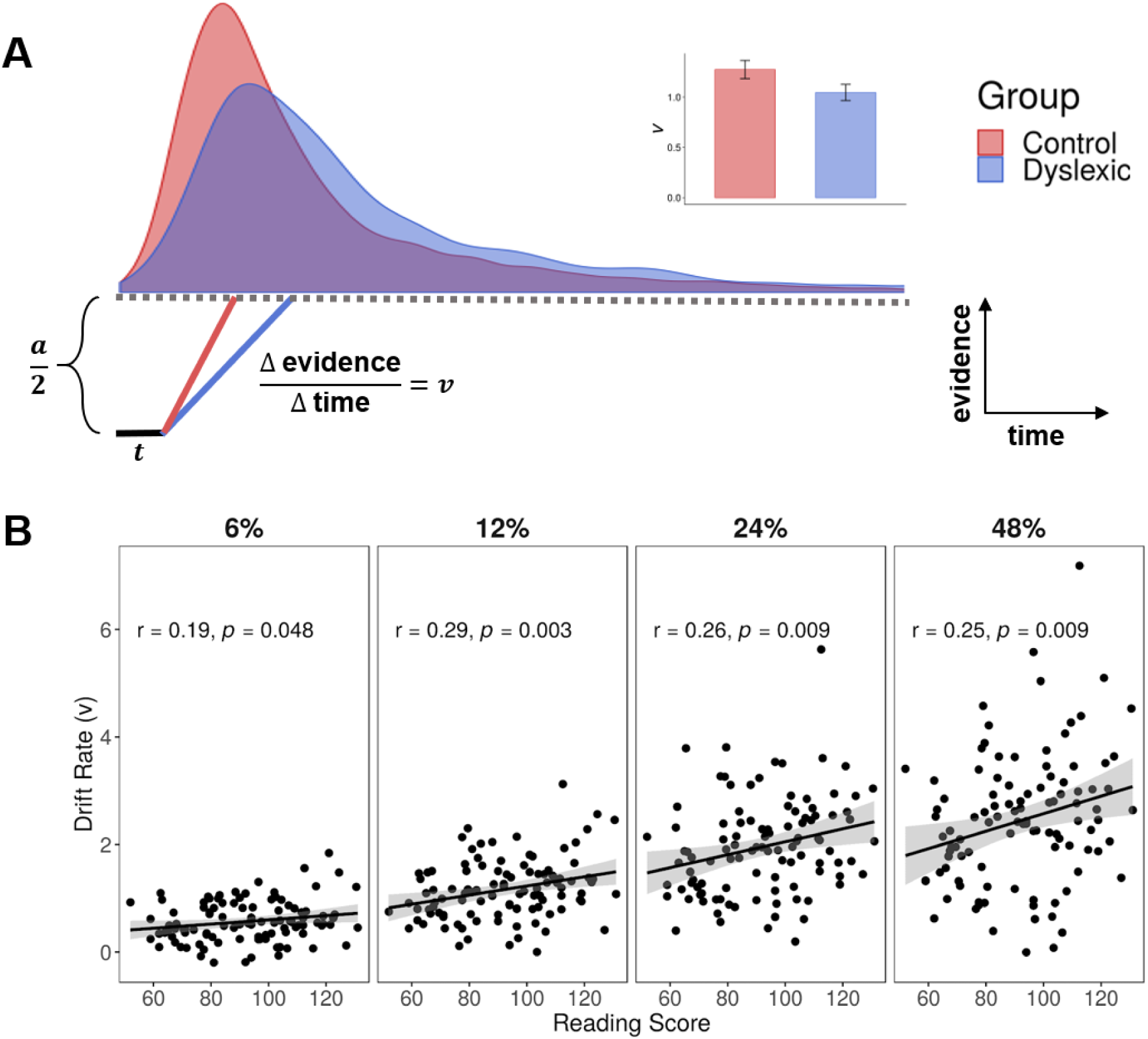
Panel (A): A schematic of the drift diffusion model (DDM) with reaction time distributions (at 12 % coherence) from the control and dyslexic groups imposed above. The red and blue lines in the schematic show how differences in drift rater predict differences in the reaction time distributions. The DDM model was fit separately to each individual’s data and the average drift rate parameter for the dyslexic and control groups is shown in the bar plot in panel A (+/− 1 standard error). (B) The relationship between estimated drift rate and reading skill at four different stimulus coherence levels. Lines are best fit regression lines and shaded regions are confidence intervals.

After fitting the DDM to each subject’s behavioral responses, we investigated whether there was a relationship between the drift rate parameter, *v*, and reading skill. Drift rate models the efficiency with which information is extracted and integrated from incoming sensory signals. For example, drift rate monotonically increases with stimulus coherence level (*β* = 0.719, SE = 0.0249, p < 1 × 10^−15^) indicating the visual system can more efficiently extract motion information when stimulus noise is low. If individuals with dyslexia do not have any difficulties with sensory encoding, as predicted by the statistical learning hypothesis, we would expect drift rate to be uncorrelated with reading skill once covariates like IQ, age, and ADHD diagnosis are controlled for. Note that in our analyses, we treat reading as a continuous measure, but we also provide analyses where reading disability is treated as a categorical variable in Supplementary Analysis 1 (Tables S7-S9).

Individual estimates of drift rate are shown in Figure 2. Drift rate was best modeled by a main effect of reading skill, a main effect of stimulus coherence, a main effect of age, and the interaction of reading skill and stimulus coherence (Table 1). Our results therefore indicate that drift rate increases with stimulus coherence, as expected, as well as age and reading skill. Furthermore, there is a stronger relationship between reading skill and drift rate at high stimulus coherence levels, which is likely a consequence of the fact that estimates of drift rate are more reliable at higher coherence levels (see Methods).

**Table 1.** Selected model of drift rate

| | $\beta$ | SE | $p$ |
| --- | --- | --- | --- |
| Intercept | 1.534 | 0.0620 | $< 1 \times 10^{-15}$ |
| Stimulus coherence | 0.719 | 0.0249 | $< 1 \times 10^{-15}$ |
| Age | 0.268 | 0.0623 | $3.878 \times 10^{-5}$ |
| Reading skill | 0.173 | 0.0623 | $6.605 \times 10^{-3}$ |
| Stimulus coherence : Reading skill | 0.0869 | 0.0249 | $7.04 \times 10^{-4}$ |

The DDM also estimates a parameter modeling the trial-to-trial variability in drift rate, *sv*. This parameter is known to be correlated with drift rate under certain conditions, with higher average drift rates being associated with greater trial-to-trial variability(Wagenmakers & Brown, 2007; Wagenmakers, Grasman, & Molenaar, 2005). Unsurprisingly, we found that *sv* was correlated with drift rates at every stimulus level. It was positively related to reading skill, but this effect did not reach significance (r = 0.13, *p =* 0.0953).

As to the question of whether drift rate explains additional variance in reading skill beyond phonological processing, consider the subset of readers in our sample with above average PA (PA scores ≥ 100). Within this subgroup of 38 participants, 9 children (23.7%) met our criteria for dyslexia despite having high PA, and reading skill was significantly correlated with mean drift rate (r = 0.49, *p =* 0.0019; see Figure 3). For these individuals, knowing drift rate explains 24% of variance in reading skill. In readers with average-or-better PA, it appears that individual differences in motion encoding and sensory integration distinguish between struggling and expert readers.

**Figure 3.**
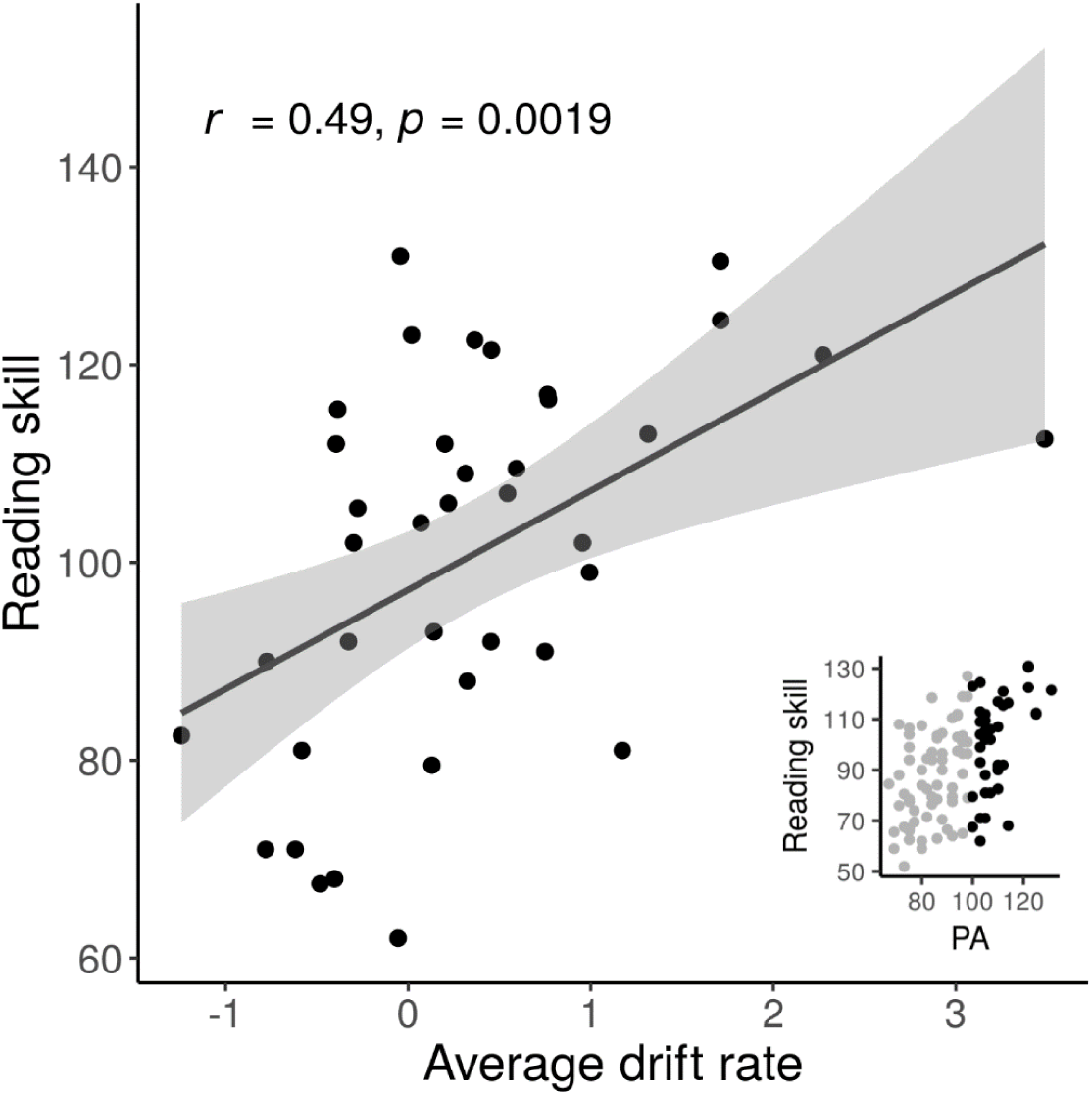
The relationship between drift rate and reading skill in a subset of individuals with good phonological awareness. Average drift rate is calculated by averaging each individual’s z-scored drift rate estimates at each stimulus coherence level. Inset: a scatter plot indicating in black which subset of the study sample is included in the “good phonological awareness” group.

### Decision making parameters are related to reading skill and independent of sensory processing

We next consider the predictions of the non-sensory hypothesis by analyzing the relationship between non-sensory parameters of the DDM and reading skill (Figure 4A-D). If poor readers struggled with the task only because of differences in sensory encoding, we would expect no parameters besides drift rate (and *sv*) to be correlated with reading skill. To the contrary, the parameter *sz* was correlated with reading skill and, after model selection, the best model of *sz* contained only a main effect of reading skill (*β* = −0.0842, SE = 0.0280, *p =* 0.00331). The parameter *sz* represents the trial-to-trial variability in the relative amount of evidence required to initiate a judgment; individuals with high *sz* values are prone to making fast errors. Indeed, we confirmed that the ratio of median correct response times to error response times within a subject was correlated with the DDM estimation of *sz* (r = 0.452, *p =* 1.44 × 10^−6^).

**Figure 4.**
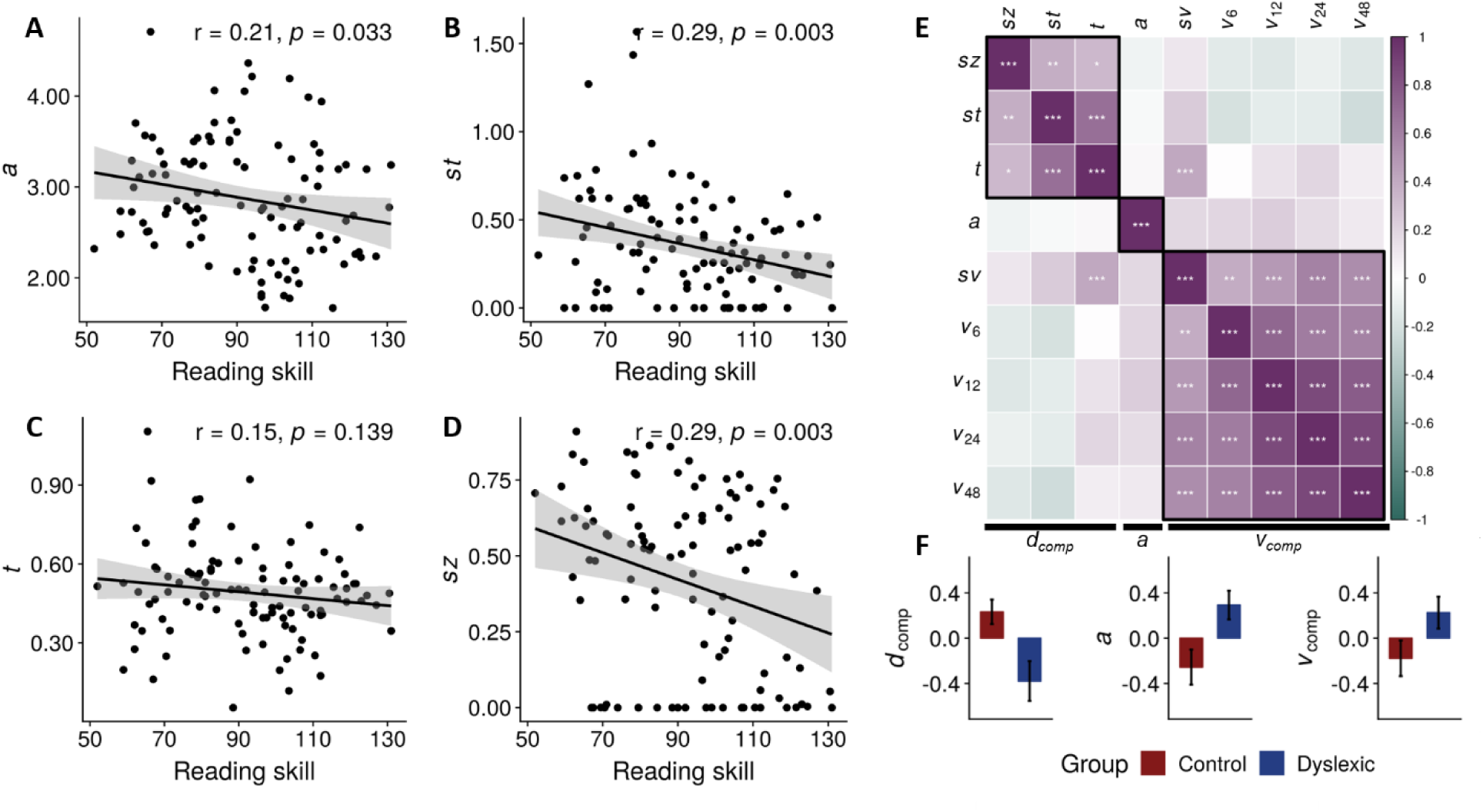
Panels A-D: The relationship between reading score and four non-sensory parameters of the DDM. (A) decision threshold a, (B) variability in drift process starting point sz, (C) non-decision time t, and (D) variability in non-decision time st. Panel E: correlations between parameters of the DDM. Boxes indicate hierarchical clustering results (Ward’s method) and stars indicate significant correlations after Holmes-Sidak correction for multiple comparisons: *p* < 0.05 is noted with *, *p* < 0.01 with **, and *p* < 0.001 with ***. Panel F: group comparisons for the three composite measures based on hierarchical clustering of the DDM parameters: *d*_comp_: composite of *sz, st*, and *t*, the a parameter, and *v*_*comp*_: composite of the four drift rate parameters and *sv*. Note that all three composite parameters are z-scored. Error bars represent one standard error of the mean.

Similarly, we observed that the parameter representing the threshold of evidence required to initiate a decision, *a*, had a modest but significant correlation with reading skill (*β* = −0.136, SE = 0.0632, *p =* 0.0329), indicating that worse reading skill is associated with employing a more conservative criterion for initiating a perceptual decision. No covariates (age, nonverbal IQ or ADHD diagnoses) were retained by model selection.

Lastly, we examined parameters that represent the lumped contributions of all non-decision processes to reaction time, including the time necessary to encode a sensory stimulus and execute a motor response. Because some individuals with dyslexia are known to have slower processing speed (Pennington et al., 2012; Peterson & Pennington, 2015), we might expect this time to be longer in children with worse reading skills. Indeed, the parameter *t* representing an individual’s average non-decision time showed an overall negative relationship with reading skill. However, the magnitude of the effect was not nearly large enough to attain statistical significance, and after model selection, only age was retained as a predictor of *t* (*β* = −0.0496, SE = 0.0163, *p =* 0.00301). As such, maturation is associated with reduced non-decision time. Interestingly, a parameter modeling trial-to-trial variability in non-decision time, *st*, was best modeled by main effects of reading skill (*β* = −0.0810, SE = 0.0278, *p =* 0.00436) and age (*β* = −0.0846, SE = 0.0278, *p =* 0.00296).

We have so far identified several parameters of the DDM, both sensory and non-sensory, that show associations with reading skill. We next considered the extent to which these parameters were correlated with one another (Figure 4E). As expected, we noted strong correlations between the four drift rate parameters. None of the drift rate parameters were significantly correlated with any non-sensory parameters after correction for multiple comparisons. There were moderate correlations between three non-sensory parameters, *st*, *t* and *sz* (*st* and *t*: r = 0.685, *p =* 9.75 × 10^−16^; *t* and *sz*: r = 0.335, *p =* 0.0005; *sz* and *st:* r = 0.386, *p =* 5.03 × 10^−5^) These three parameters largely contribute to modeling the leading edge of the reaction time distribution: *sz* allows for the presence of relatively fast errors, *t* shifts the response time distribution along the time axis, and *st* allows for responses before an individual’s average response time. Finally, we noted that the parameter *a* was uncorrelated with any of the other parameters.

Hierarchical clustering with Ward’s method (Ward, 1963) indicated that the correlation matrix was consistent with three clusters of parameters: a cluster consisting only of *a*, another consisting of the *st*, *t*, and *sz*, and a final cluster including all four drift rates and *sv*. This suggests that the DDM captures several independent mechanisms underlying sensory encoding and perceptual decision making.

### Sensory and non-sensory predictors both explain reading outcomes

So far in our analysis, there seem to be several separate profiles of performance on the motion discrimination task that are associated with low reading skill: reduced ability to encode and integrate sensory information, setting a more conservative decision criterion, and generally more variability in terms of the time taken to gather evidence and/or execute a decision. The lack of correlations between many of the DDM parameter estimates indicates that individuals who display a deficit in terms of one process (e.g., sensory encoding), are not necessarily the same individuals who perform abnormally in terms of another process (e.g., decision-making) and that profiles of performance are variable across subjects. Therefore, we might expect that each parameter contributes separately to explaining variance in reading outcomes.

To test whether each dimension of task performance is indeed a unique contributor to a model of reading skill, we employed a linear model. To simplify the number of parameters, we introduce several composite measures based on the correlation matrix of DDM parameters and our clustering analysis (Figure 4F). Drift rate is summarized as a composite measure, *v*_*comp*_, by taking the first principal component of the four drift rates and *sv*. A second composite measure *d*_*comp*_ was derived from the first principal component *st*, *t*, and *sz*, which we expect represents aspects of variability in the decision-making process.

The dyslexic and control groups differed in terms of each of these three mechanisms (Figure 4F). We performed model selection, starting with the full model with reading score as the dependent measure and all hypothesized DDM parameters and the three covariates (*v*_*comp*_, *d*_*comp*_, *a*, nonverbal IQ, ADHD diagnosis and age) as predictors. The selected model retained all three predictors from the DDM and nonverbal IQ (Table 2).

**Table 2.**
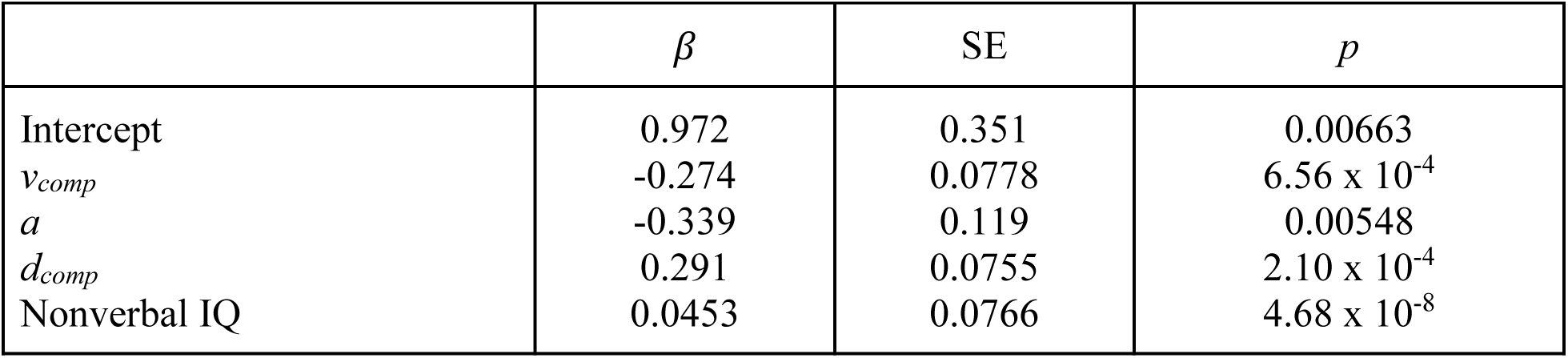
Selected model of reading skill from DDM parameters

This result confirms that non-sensory mechanisms explain additional variance in reading skill once the quality of sensory evidence encoding is accounted for. As such, even within this single psychophysical task, there are multiple non-correlated dimensions of variance contributing to the pattern of responses observed in individual’s with dyslexia: the ability to extract evidence from sensory information, choice of decision threshold, and trial-to-trial variability in behavior.

### Psychophysical measures are not proxies for PA

To address the question of whether performance on the motion discrimination task is related to reading skill by way of phonological processing, or in addition to it, we explore a series of models. We first test the hypothesis that predictors from the psychophysical task do not explain additional variance in reading skill once phonological processing is accounted for. We again modeled reading skill as a function of our parameters of interest from the DDM—*v*_*comp*_, *d*_*comp*_, and *a*—as well as two phonological processing measures, PA and RAN, and the three covariates. Model selection retained all predictors except ADHD diagnosis and age (Table 3). Correspondingly, an ANOVA F-test comparing the selected model to a reduced model with only PA, RAN and nonverbal IQ confirmed that adding predictors from the DDM explained variance in reading skill above and beyond the reduced model (F(100, 97) = 4.0438, *p =* 0.00936). The reduced model also had a higher AIC (selected model AIC = 794.4, reduced model AIC = 800.7) and BIC (selected model BIC = 813.9, reduced model BIC = 815.6). From this analysis, we can confirm that all three predictors from the DDM are useful for explaining differences in reading skill *in addition* to traditional measures of phonological processing.

**Table 3.**
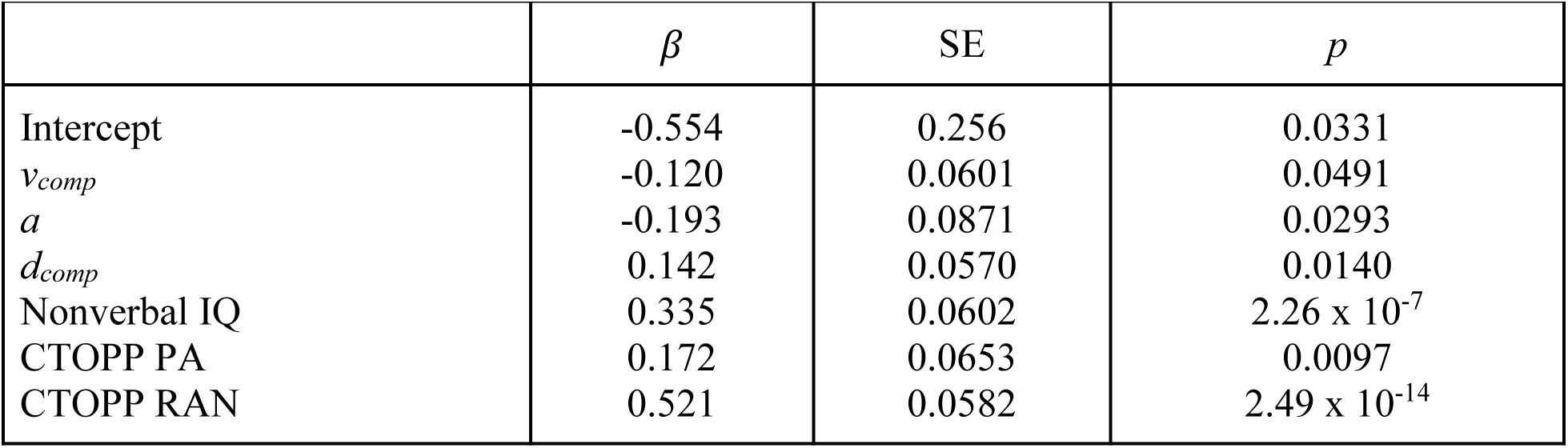
Selected model of reading skill

| | $\beta$ | SE | $p$ |
| --- | --- | --- | --- |
| Intercept | -0.554 | 0.256 | 0.0331 |
| $v_{comp}$ | -0.120 | 0.0601 | 0.0491 |
| $a$ | -0.193 | 0.0871 | 0.0293 |
| $d_{comp}$ | 0.142 | 0.0570 | 0.0140 |
| Nonverbal IQ | 0.335 | 0.0602 | $2.26 \times 10^{-7}$ |
| CTOPP PA | 0.172 | 0.0653 | 0.0097 |
| CTOPP RAN | 0.521 | 0.0582 | $2.49 \times 10^{-14}$ |

Because ordinary least squares models may be poorly affected by multicollinearity, we also applied lasso regression with 10-fold cross validation (Friedman, Hastie, & Tibshirani, 2010). This modeling approach is provided in the Supplement (Figures S3-S4; Table S10).

### Do sensory deficits have cascading effects?

It has been argued that deficits in sensory processing or decision-making could affect reading skill because they disrupt the typical development of PA (Lieder et al., 2019; Manis et al., 1997; Richardson, Thomson, Scott, & Goswami, 2004). We therefore explored whether this hypothesis is borne out in our data by employing a mediation analysis. While *a* was not significantly correlated with PA, *v*_*comp*_ and *d*_*comp*_ showed modest correlations (*v*_*comp*_ and PA: r = 0.324, *p =* 4.80 × 10^−4^; *d*_*comp*_ and PA: r = 0.182, *p =* 0.0358).

We first tested a model with PA mediating the relationship between *v*_*comp*_ and reading skill and found a significant, partial mediation effect (42.3%, *p =* 0.0052). Similarly, the *d*_*comp*_-reading skill relationship is partially mediated by PA (22.2% mediation, *p =* 0.0224)), but there was also still a significant direct relationship (*β* = 4.293, SE = 1.501, *p =* 0.00516). As such, our results provide some support for the idea that in certain poor readers, low PA could be a consequence of a more fundamental impairment in either sensory or non-sensory mechanisms. However, our data suggest a partial mediation, indicating that many individuals would not be well described by this cascading model and that there are also direct links between the model parameters and reading skill.

### Multiple dimensions of skilled and disabled reading

Contrary to theories that seek to discover a unified deficit that characterizes children with dyslexia, we have established that sensory processing of visual motion is separable from non-sensory aspects of perceptual decision making, and both factors account for independent variance in reading skill. To speak to the question of how many separable underlying factors predict reading skill, we next apply exploratory factor analysis (EFA). EFA is an unsupervised learning approach for identifying the number, and characteristics, of *latent factors* that explain the correlation structure of a multi-dimensional data set (Costello & Osborne, 2005; Ferguson & Cox, 1993; Kline, 2013). We applied EFA to characterize the space of the DDM parameters, nonverbal-IQ, and the six subtests of the CTOPP (measure of reading skill were not included in the EFA). An analysis of the eigenvalues of the correlation matrix indicated that four latent factors were warranted (i.e., the first four eigenvalues > 1, see scree plot in Figure S5). This was confirmed by parallel analysis (Hayton, Allen, & Scarpello, 2004) (i.e., in a simulation of 1000 random correlation matrices, the first four resulting eigenvalues were lower than the corresponding eigenvalues from our data’s correlation matrix 95% of the time). The four factors are shown in Figure 5 with orthogonal varimax rotation. The total proportion of explained common variance by the four-factor model was 55.8% (Factor 1: 20.3%, Factor 2: 14.2%, Factor 3: 10.7%, Factor 4: 10.6%).

**Figure 5.**
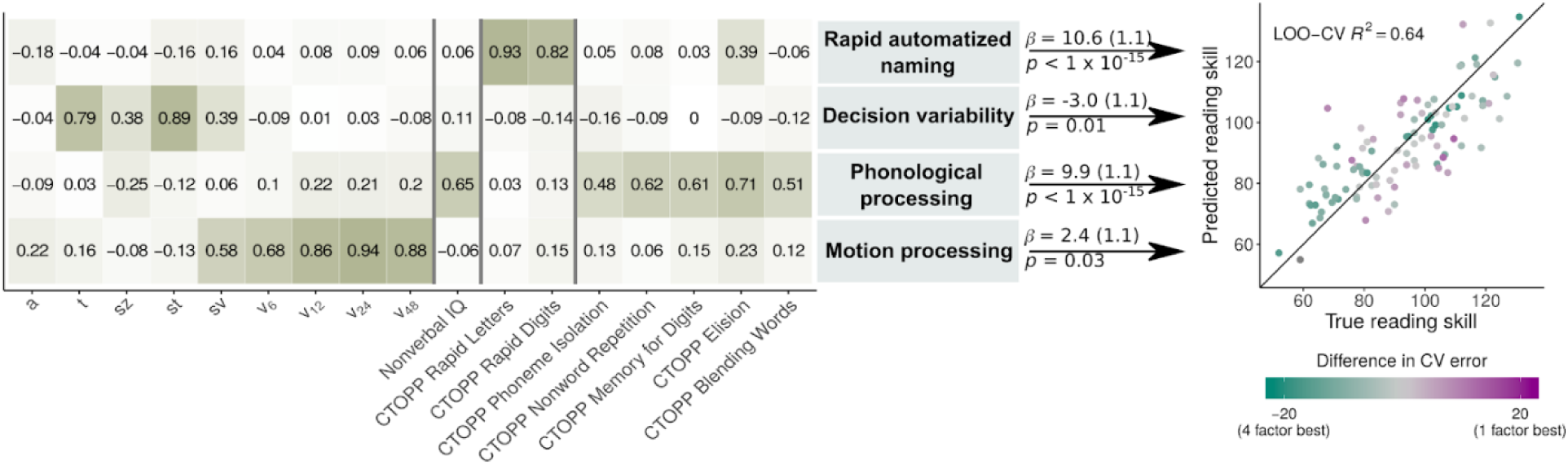
Factor loadings for the orthogonal four-factor model are shown in the table; shading corresponds to absolute value of the loading. The scatterplot shows the correspondence between true (measured) and predicted reading skill using a linear model with all four factors as predictors. Each point was predicted using leave-one-out cross-validation (LOO-CV). Color indicates whether that point was more accurately predicted by the single-factor model or the full model with all four factors. Green points had a lower squared error when predicted by the four-factor model, and purple points had a lower squared error when predicted by the single-factor. Gray points had similar prediction accuracy for both models.

The loadings of the first factor are dominated by the four drift rate parameters, whereas the second factor is loaded most heavily by nonverbal-IQ and four of the CTOPP subtests. The remaining two subtests, Rapid Digits and Rapid Letters, load onto their own factor (in line with the double-deficit hypothesis (Wolf & Bowers, 1999)). An additional factor appears to reflect non-decision time and variability parameters of the DDM *st*, *sz*, and *t*. Notably, the evidence threshold parameter, *a*, is not particularly associated with any factor; 87% of variance in *a* is unexplained by this model.

Factor analysis largely conforms to the intuitions we have built so far through linear models: drift rate, although correlated with phonological processing and perhaps partially mediated by it, is identified as a separate factor. Drift rate and the non-sensory parameters of the DDM are modeled as observations from two distinct factors. It is likely that *a* is representative of an additional factor, consistent with its lack of correlations with any other parameter of the DDM (note that without multiple estimates of *a*, EFA cannot estimate measurement noise and consequently does not assign it to a new factor). Critically, each of these four factors was significantly related to reading skill demonstrating that, rather than representing a single underlying construct, there are multiple, independent cognitive and sensory dimensions characterizing individual differences in reading skill (Figure 5). A linear model of reading skill as a function of scores on the four factors indicated that all four effects were significant (see coefficients in Figure 5). Furthermore, the full model also had a lower AIC (full model AIC = 798.8, single factor model AIC = 869.9) and BIC (full model BIC = 814.6, single factor model BIC = 877.8).

In addition to standard model selection, we compared the accuracy of the four-factor model on predicting held-out observations to the accuracy of a single-factor model. Using leave-one-out cross validation to control for overfitting, the four-factor model explained 63.9% of variance in reading skill for the held-out points. The single factor model used only Factor 2, which is largely a composite of the CTOPP measures of PA, phonological memory, and nonverbal IQ. This model was only able to explain 27.4% of variance in reading skill for held-out observations (Figure S6), indicating the necessity of considering multiple underlying dimensions (at least 4) in order to accurately predict individual differences in reading ability.

## Conclusions

Our results demonstrate that (1) a core phonological deficit model is insufficient to account for many cases of developmental dyslexia, (2) abnormal performance on the motion discrimination experiment in children with dyslexia cannot be ascribed to a uniform profile of either sensory or non-sensory deficits, (3) both sensory and non-sensory mechanisms explain variance in reading skill above and beyond phonological processing, (4) the correlational structure of cognitive, linguistic and sensory measures explored here is consistent with, at minimum, four underlying factors, (5) each of these four factors accounts for unique variance in children’s reading abilities. In sum, our results are not consistent with models of dyslexia that only consider phonological processing or models in which impairments in sensory encoding or decision making primarily affect reading development via a disruption of phonological processing. Instead, dyslexia should be conceptualized as a disorder that may arise from several distinct loci.

Our work is consistent with that of the Pennington and colleagues, which has capitalized on large samples to demonstrate that individuals with dyslexia have a heterogeneous profile of cognitive and linguistic impairments (Pennington, 2006; Pennington et al., 2012; Peterson & Pennington, 2015). The present work extends this perspective to address the role of sensory processing and perceptual decision-making deficits in dyslexia.

Several preceding studies have attempted to investigate multiple candidate mechanisms of dyslexia, including auditory, visual, and motor processes. Our work generally conforms to the finding of at least four such studies (Ho et al., 2002; Menghini et al., 2010; Ramus et al., 2003; White et al., 2006) that show a heterogenous pattern of deficits present in struggling readers. In a study with related methodology, Talcott *et al.* collected several psychophysical measures in 350 school aged children and found that each uniquely explained a small percentage of variance in literacy skill (Talcott et al., 2000). Our study similarly finds that several distinct mechanisms each explain a small, but unique, proportion of variance in reading outcomes.

To our knowledge, the present study is the first use of the DDM to model motion discrimination in children with dyslexia. Our results serve as a partial validation of two seemingly contradictory theories: some poor readers show a pattern of performance consistent with reduced ability to extract information from incoming sensory signals, while others are better described as having normal sensory processing but altered decision-making characteristics (including, as the propensity to make fast errors reveals, more trial-to-trial variability in the relative amount of evidence needed to initiate a decision). Neither the statistical learning hypothesis, which would argue that sensory deficits are not meaningful, nor the magnocellular deficit hypothesis, which would fail to predict the non-sensory parameters of the DDM relate to reading skill, entirely match our results. Yet we see evidence for both sensory- and non-sensory profiles of impairment in our sample. In line with these findings, we propose that each mechanism should be reconceptualized as a dimension of risk, as opposed to a single cause, of reading difficulties.

As a correlational study, our results cannot validate any particular causal mechanism. It is possible that each factor represent clusters of symptoms that indicate underlying impairment in a processing system, but are not a direct cause of dyslexia themselves. For example, the fact that differences in visual motion processing predict unique variance in reading skill does not necessarily mean that, for those individuals, poor perception of visual motion is the cause of their reading difficulty. Instead, measurements of task performance may be a proxy for the fidelity with which the visual system constructs a sensory representation of a noisy stimulus (Sperling, Lu, Manis, & Seidenberg, 2005, 2006), or the efficiency of information transfer between visual regions (Yeatman, Dougherty, Ben-Shachar, & Wandell, 2012; Yeatman, Rauschecker, & Wandell, 2013), or the integration of sensory signals over time (Joo et al., 2017). Skilled reading requires rapid communication among a distributed network of visual, auditory and language processing systems and an impairment in any one of these systems, or the connections between them, could cause difficulties learning a complex skill like reading (Wandell & Yeatman, 2013).

Our main conclusion is a lack of concordance with either a single deficit or cascading model. As such, our results contradict claims that a single mechanism, either phonological or sensory, can be considered the “fundamental” or “core” deficit of dyslexia. In particular, our work opposes the recent claim that the majority of individuals with dyslexia have a magnocellular processing deficit (Stein, 2018); if the DDM is accepted as a reasonable model of behavior on the motion discrimination task—a starting point with considerable basis (Huang-Pollock et al., 2017; Palmer et al., 2005; Ratcliff & McKoon, 2008)—then we conclude that a minority of children with dyslexia are best modeled as having a motion encoding deficit.

Furthermore, while we do not directly test auditory theories of dyslexia here, our results still speak to this research. For example, the influential temporal sampling hypothesis holds that the core deficit of dyslexia is abnormal processing of syllable-scale acoustic features, which in turn disrupts PA development and manifests as sampling problems in the visual domain (Casini et al., 2018; Goswami, 2011). Our results indicate that, even if we could establish that abnormal auditory processing impairs PA, many cases of dyslexia would still be unaccounted for based on the effectiveness of a phonological-core model. Furthermore, the idea that difficulties sampling incoming stimuli largely explains poor performance on the motion discrimination task is specious, as we have demonstrated that there are several reasons (some non-sensory) why individuals with dyslexia may perform differently on this task than typical readers. While the idea of a centralized deficit in some aspect of temporal processing has an elegant appeal, our data are simply not consistent with such a simple model.

The clinical implications of this multifactorial model are a target for future research. Whether or not different risk profiles predict outcomes for children enrolled in competing intervention programs is an empirical question that cannot be readily inferred from correlational data. For example, in a previous intervention study we demonstrated that individual differences in visual motion sensitivity have no prognostic value for predicting a child’s response to intervention (Joo et al., 2017).

Moving forward, we propose an additive risk factor mode of dyslexia in which multiple dimensions of sensory, cognitive and linguistic processes contribute distinct risk for reading difficulties. Our results are agnostic to whether poor performance on any given task indicates deficits in the specific targeted function (e.g., motion processing) or indexes processing capacities of a broader system (e.g., constructing a high-fidelity representation of a noisy visual signal). There are many proposed neurobiological mechanisms that could, in theory, be compatible with our findings (e.g., heterogeneous profiles of abnormal cortical migration (Hancock, Pugh, & Hoeft, 2017)).

In sum, our results demonstrate that an additive model outperforms cascading deficit models or models that only consider measures of phonological processing without considering the role of sensory processing and perceptual decision making. Thus, rather than continuing to seek an underlying cause of dyslexia, the field should systematically build towards a more complete model of the factors that add risk (or protection) for reading difficulties. Our data and model necessitate a shift towards theories that explain skilled and disabled reading as emerging from a high-dimensional space determined by several distinct processing systems.

## Acknowledgements

We would like to thank Sung Jun Joo for helpful discussions that influenced our study design and psychophysics protocol. G.O. was supported by the Auditory Neuroscience Training Grant (NIH grant T32DC005361-16). This work was also supported by the National Science Foundation, Division of Behavioral and Cognitive Sciences Grant 1551330, Eunice Kennedy Shriver National Institute of Child Health and Human Development Grant P50 HD052120 and R21 HD092771 and research grants from Microsoft to J.D.Y.

