## Supplemental Material for "Bridging sensory and language theories of dyslexia: towards a multifactorial model"

Supplementary Materials

| Table S1: Correlations with composite reading score | | | |
| --- | --- | --- | --- |
| Measure | 𝛽 | SE | *P* |
| Age  ADHD diagnosis  Gender | 1.08  -6.91  1.01 | 1.4  4.45  3.78 | 0.443  0.124  0.789 |
| **CTOPP-2** | | | |
| Phonological Awareness  Phonological Memory  Rapid Automatic Naming | 0.65  0.57  0.88 | 0.11  0.1  0.09 | 3.46 x 10^-8^  8.10 x 10^-8^  1.23 x 10^-15^ |
| **TOWRE-2** | | | |
| Pseudoword Decoding  Sight Word Reading | 1.02  0.82 | 0.03  0.03 | < 1 x 10^-15^  < 1 x 10^-15^ |
| **WASI-II** | | | |
| Full-scale IQ  Matrix Reasoning t-score  Vocabulary t-score | 0.75  0.95  0.97 | 0.09  0.16  0.13 | 2.87 x 10^-13^  3.50 x 10^-8^  4.05 x 10^-11^ |
| **Woodcock Johnson** | | | |
| Letter Word Identification  Word Attack | 0.91  0.94 | 0.03  0.05 | < 1 x 10^-15^  < 1 x 10^-15^ |

| Table S2: Group demographic information | | | | | |
| --- | --- | --- | --- | --- | --- |
|  | **Mean** | | **SD** | | *p* |
|  | Control  *n* = 48 | Dyslexic  *n* = 43 | Control | Dyslexic |  |
| Age  ADHD diagnosis  Gender (♂) | 10  4  28 | 9.69  13  25 | 1.38 | 1.15 | 0.245  0.009  0.985 |
| **CTOPP-2** | | | | | |
| Phonological Awareness  Phonological Memory  Rapid Automatic Naming | 98.33  98.58  99.42 | 86.42  84.86  79 | 14.98  17.32  12.45 | 12.72  12.75  9.63 | 9.09 x 10^-5^  3.96 x 10^-5^  1.15 x 10^-13^ |
| **TOWRE-2** | | | | | |
| TOWRE Index  Pseudoword Decoding  Sight Word Efficiency | 106.79  104.6  108.17 | 68.35  71.58  68.47 | 11  11.38  11.66 | 8.01  6.32  11.3 | < 1 x 10^-15^  < 1 x 10^-15^  < 1 x 10^-15^ |
| **WASI-II** | | | | | |
| Full-Scale IQ  Matrix Reasoning t-score  Vocabulary t-score | 115.77  55.33  63.15 | 96.28  46.63  49.14 | 16.02  10.75  10.99 | 9.87  7.28  7.82 | 5.53 x 10^-10^  1.72 x 10^-5^  4.36 x 10^-10^ |
| **Woodcock Johnson** | | | | | |
| Basic Reading Score  Letter Word Identification  Word Attack | 110.27  109.75  109.35 | 77.23  74.37  82.4 | 12.71  11.79  14.89 | 10.45  12.12  10.84 | < 1 x 10^-15^  < 1 x 10^-15^  6.34 x 10^-16^ |

| Table S3: DDM parameter reliability estimates | | |
| --- | --- | --- |
| Parameter | Split-half reliability^1^ | Adjusted reliability^2^ |
| *v_6_*  *v_12_*  *v_24_*  *v_48_*  *a*  *t*  *sv*  *sz*  *st* | 0.093  0.397  0.353  0.512  0.531  0.594  0.141  0.279  0.470 | 0.170  0.568  0.522  0.677  0.693  0.737  0.248  0.436  0.640 |

^1^ Split half reliability is calculated by partitioning each subject’s responses into two by random assignment, estimating the DDM parameters on each half, and measuring the Pearson’s correlation between parameter estimates.

^2^ Adjusted reliability is calculated with the Spearman-Brown prophecy formula. This adjustment is an estimate of the reliability had the estimates been computed on twice as many observations as in split-half reliability.

| Table S4: Selected model of reaction time on the motion discrimination task | | | |
| --- | --- | --- | --- |
|  | 𝛽 | SE | *p* |
| Intercept  Stimulus coherence  Age  Reading skill | 4.101  -0.173  -0.0590  -0.00600 | 0.316  0.00898  0.0214  0.00149 | < 1 x 10^-15^  < 1 x 10^-15^  0.00703  0.000115 |

| Table S5: Selected model of accuracy on the motion discrimination task | | | |
| --- | --- | --- | --- |
|  | 𝛽 | SE | *p* |
| Intercept  Stimulus coherence  Age | 0.218  0.0844  0.0244 | 0.0618  0.00284  0.00608 | 0.000600  < 1 x 10^-15^  0.000109 |

| Table S6: Selected model of the ratio of error-to-correct-response-times  on the motion discrimination task | | | |
| --- | --- | --- | --- |
|  | 𝛽 | SE | *p* |
| Intercept  Reading skill  Age  Nonverbal IQ | 0.555  0.004440  0.08029  -0.007 | 0.3495  0.002235  0.027793  0.004208 | 0.11534  0.04966  0.00472  0.09859 |

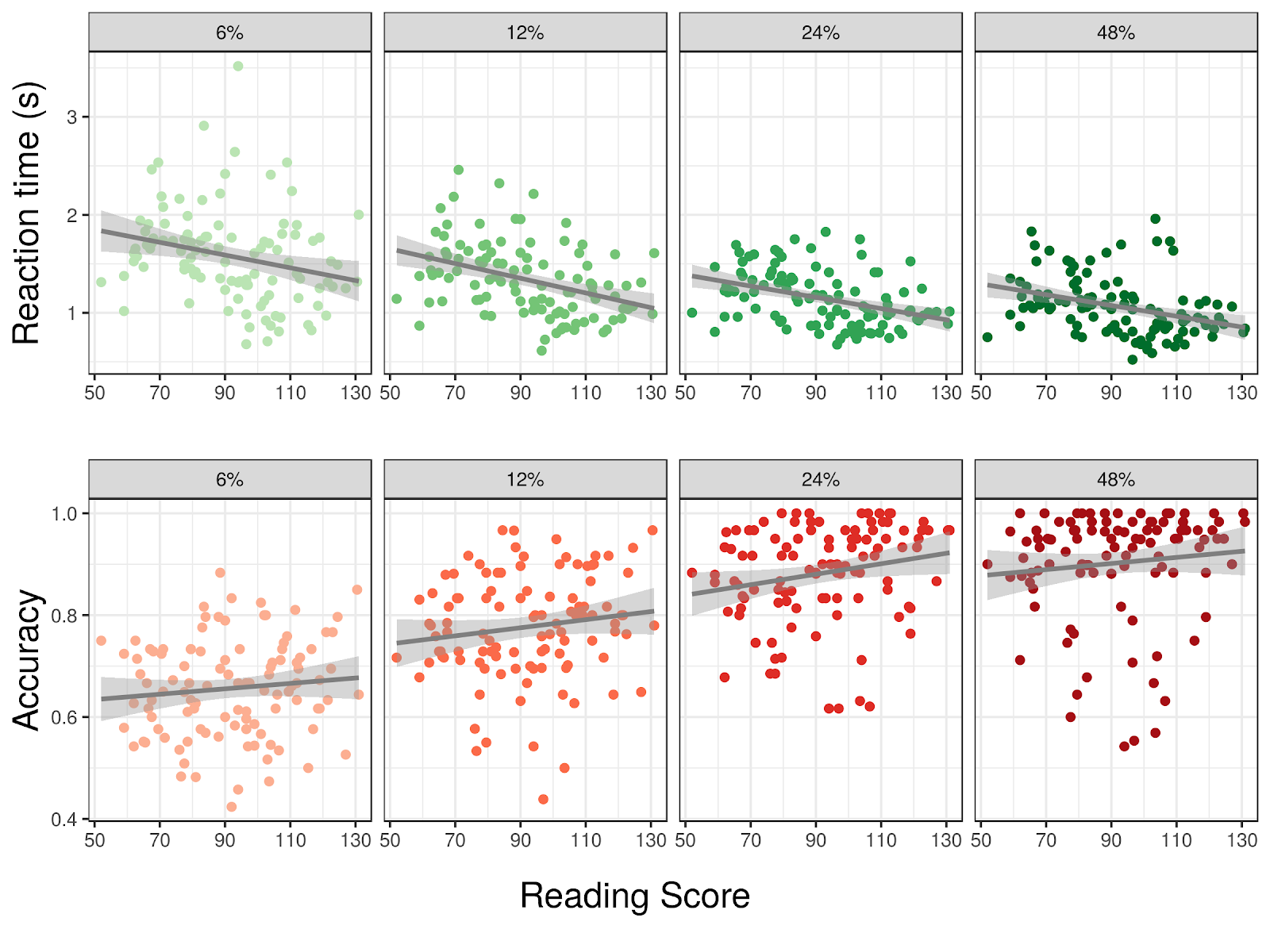

**Figure S1.** *Median reaction times (top row) and accuracy (bottom row) for each individual as a function of reading score. Panels show each stimulus coherence.*

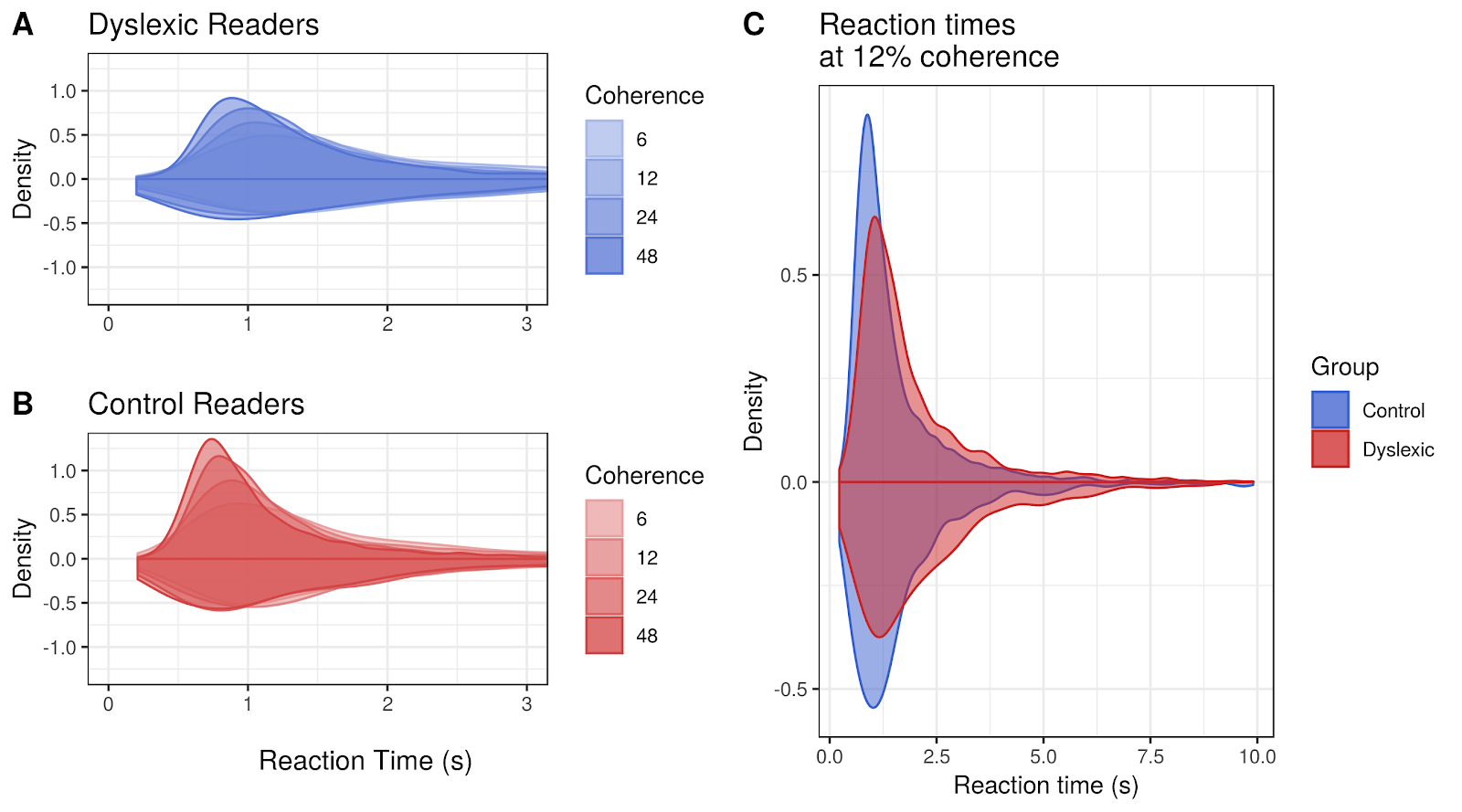

**Figure S2.** *Panels A-B: density plots of reaction times at each coherence level for the Dyslexic and Control groups. On the positive axis, correct response time distributions are shown, and on the negative axis error responses are shown. Plots are truncated at 3 seconds for ease of viewing. Panel C: overlaid density plots of reaction times at 12% coherence for the Dyslexic and Control groups.*

**Supplemental Analysis 1: group level statistics**

*Drift rate:* As in the main manuscript, we used mixed model selection to identify the most parsimonious model of drift rate. Subject was included as a random effect, as our data contained four drift rate estimates per participant (one at each coherence level). The selected model included significant main effects of stimulus coherence and age, plus a significant interaction of stimulus coherence and group (Table 4). The main effect of group was not significant (p = 0.117), but the direction of the relationship was the same as in the model where reading is treated as a continuous variable. Therefore, our results from both models are in qualitative agreement. The fact that reading skill was a significant main effect as a continuous measure but not as a discrete one is likely the result of reduced statistical power: there are 90 subjects in the group analysis, but 104 in the continuous-measure analysis.

| Table S7: Selected model of drift rate | | | |
| --- | --- | --- | --- |
|  | 𝛽 | SE | *p* |
| Intercept  Stimulus coherence  Age  Group  Stimulus coherence : Group | 1.630  0.772  0.240  -0.227  -0.133 | 0.0973  0.0265  0.0716  0.143  0.0529 | <2e-16  <2e-16  0.00117  0.117  0.0136 |

*Decision criterion parameters*: The parameter *a*, representing an individual’s threshold of evidence for initiating a decision, was modeled as the dependent variable next. Linear mixed model selection dropped all three covariates, leaving only a main effect of group (𝛽 = 0.351, SE = 0.130, p = 0.00858).

The parameter *sz*, representing trial-to-trial variability in the drift process starting point, was modeled similarly. The selected model contained only a main effect of group, although this effect missed the standard threshold of significance (𝛽 = 0.105, SE = 0.06035, p = 0.0845).

*Non-decision time parameters:* The selected model for residual non-decision time *t* contained three main effects: group, nonverbal IQ and age, of which only group fell below the standard threshold of significance (Table 5).

| Table S8: Selected model of residual time *t* | | | |
| --- | --- | --- | --- |
|  | 𝛽 | SE | *p* |
| Intercept  Group  Nonverbal IQ  Age | 0.459  0.0921  0.0321  -0.0313 | 0.0253  0.0393  0.0195  0.0179 | <2e-16  0.0214  0.1035  0.0845 |

Lastly, we considered the trial-to-trial variability in non-decision time *st*. The selected model contained two predictors: group and age at testing (Table S6).

| Table S9: Selected model of trial-to-trial variability in residual time s*t* | | | |
| --- | --- | --- | --- |
|  | 𝛽 | SE | *p* |
| Intercept  Group  Age | 0.280  0.185  -0.0644 | 0.0422  0.0622  0.0311 | 2.77e-9  0.0037  0.0410 |

We can therefore see that the relationships between estimated DDM parameters and reading skill are qualitatively consistent regardless of whether reading disability is treated as a categorical or continuous variable.

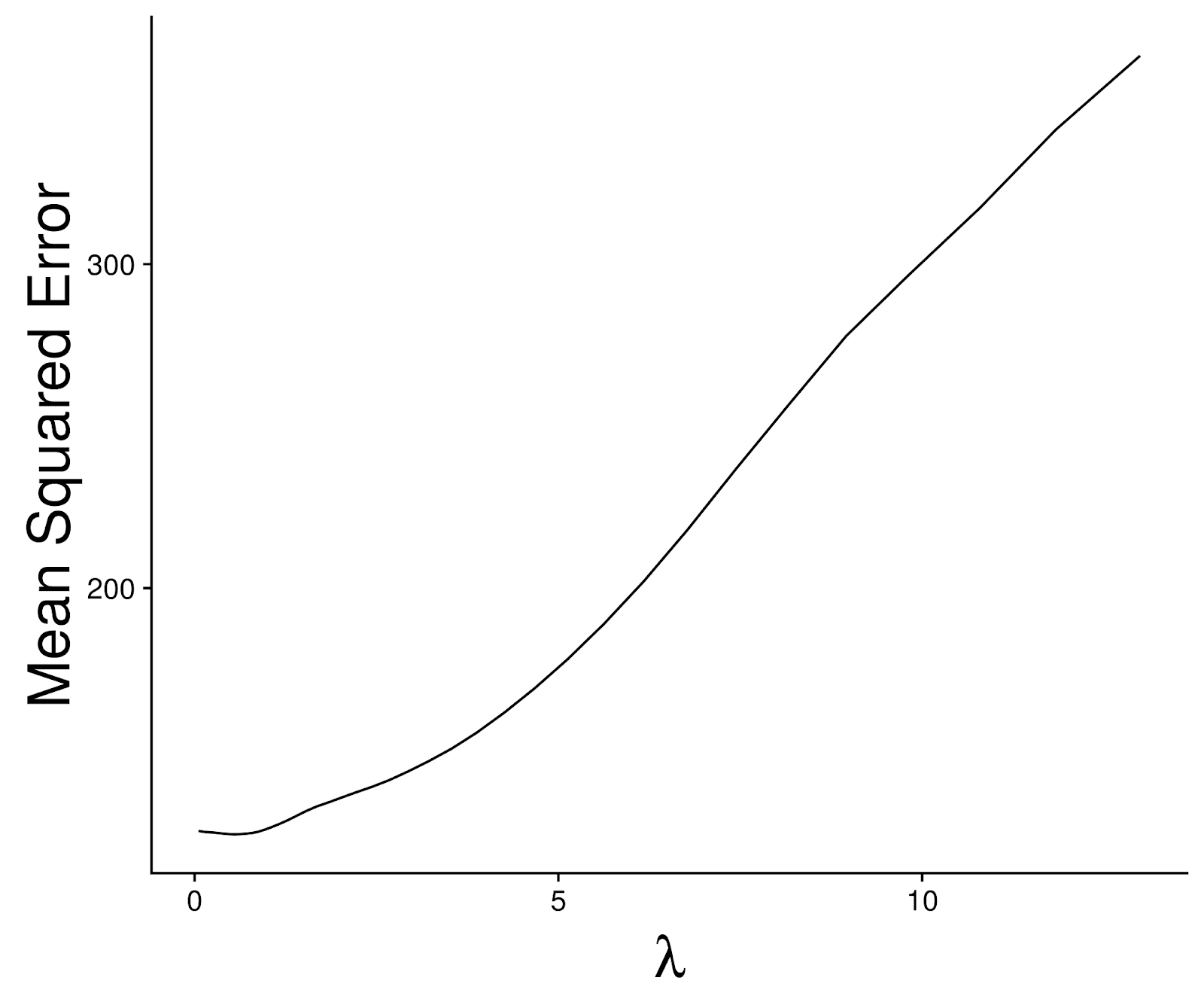

**Figure S3.** *Mean squared error of lasso regression as a function of the regularization parameter λ with 10-fold cross validation.*

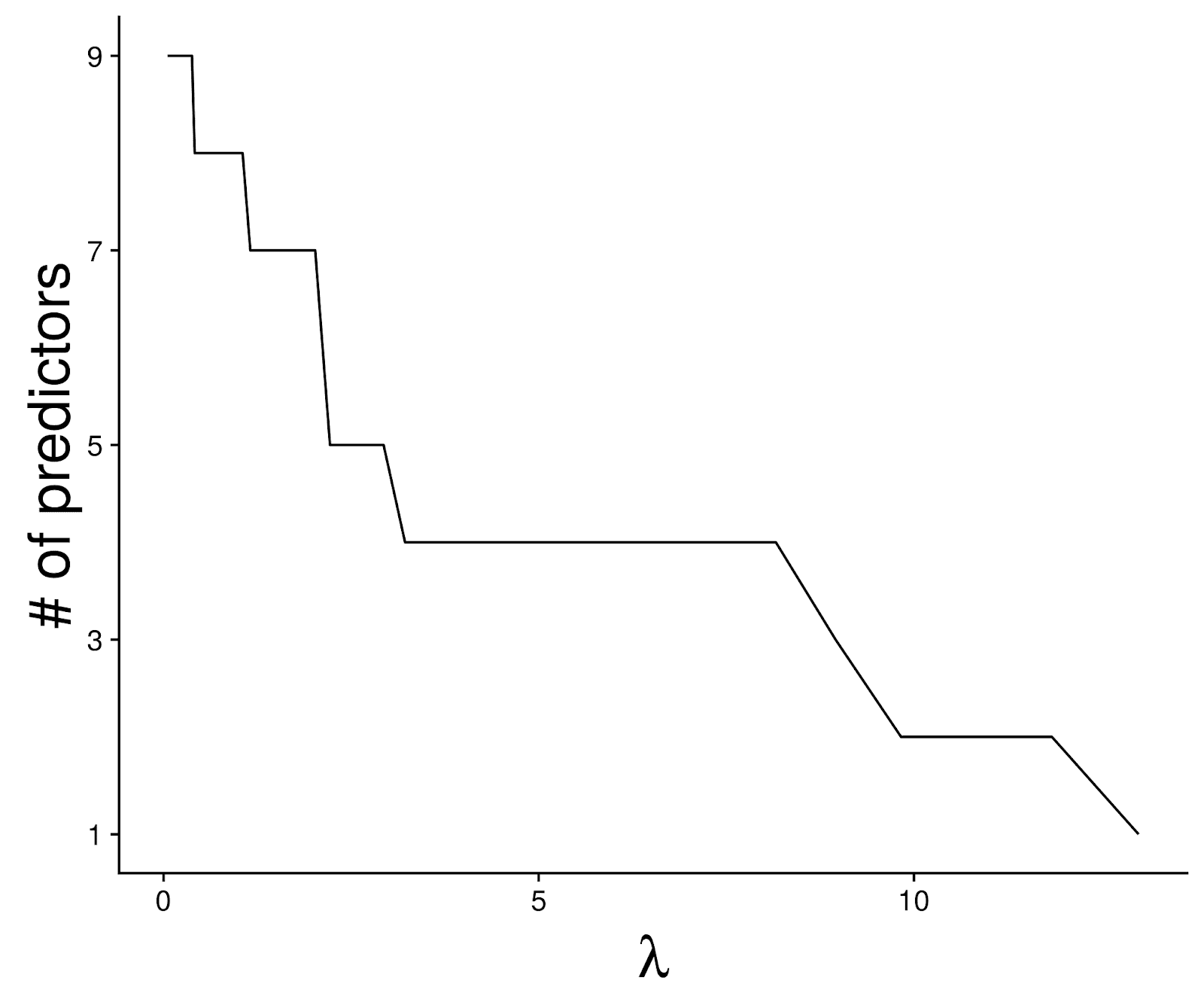

**Figure S4.** *Number of predictors retained by lasso regression as a function of the regularization parameter λ with 10-fold cross validation.*

| Table S10: Lasso model of reading skill | |
| --- | --- |
|  | 𝛽  *(*all measures are scaled prior to lasso regression)* |
| Intercept  *a*  *v_comp_*  *d_comp_*  Nonverbal IQ  CTOPP RAN  CTOPP PA  ADHD diagnosis | -3.277 x 10^-16^  -0.09240  -0.0895  0.118  0.317  0.512  0.169  -0.00221 |

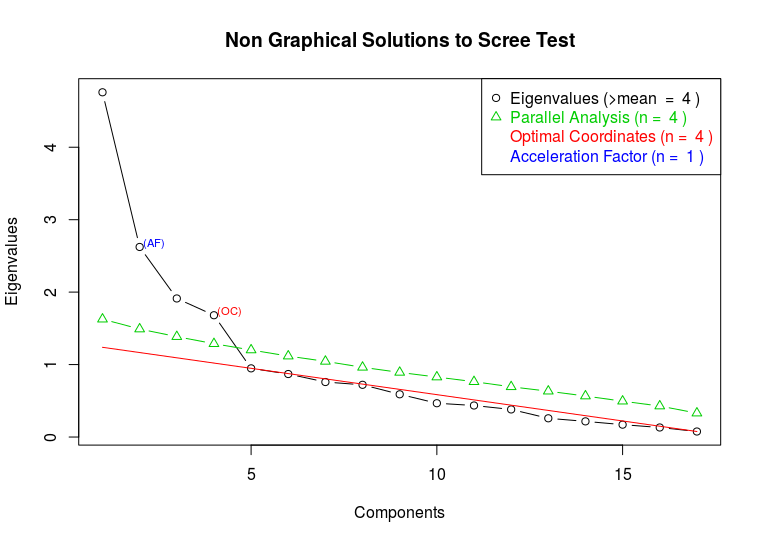

**Figure S5.** *Scree test for exploratory factor analysis. Four standard measures of model fit are given: eigenvalues, parallel analysis, the optimal coordinates metric and acceleration factor metric.*

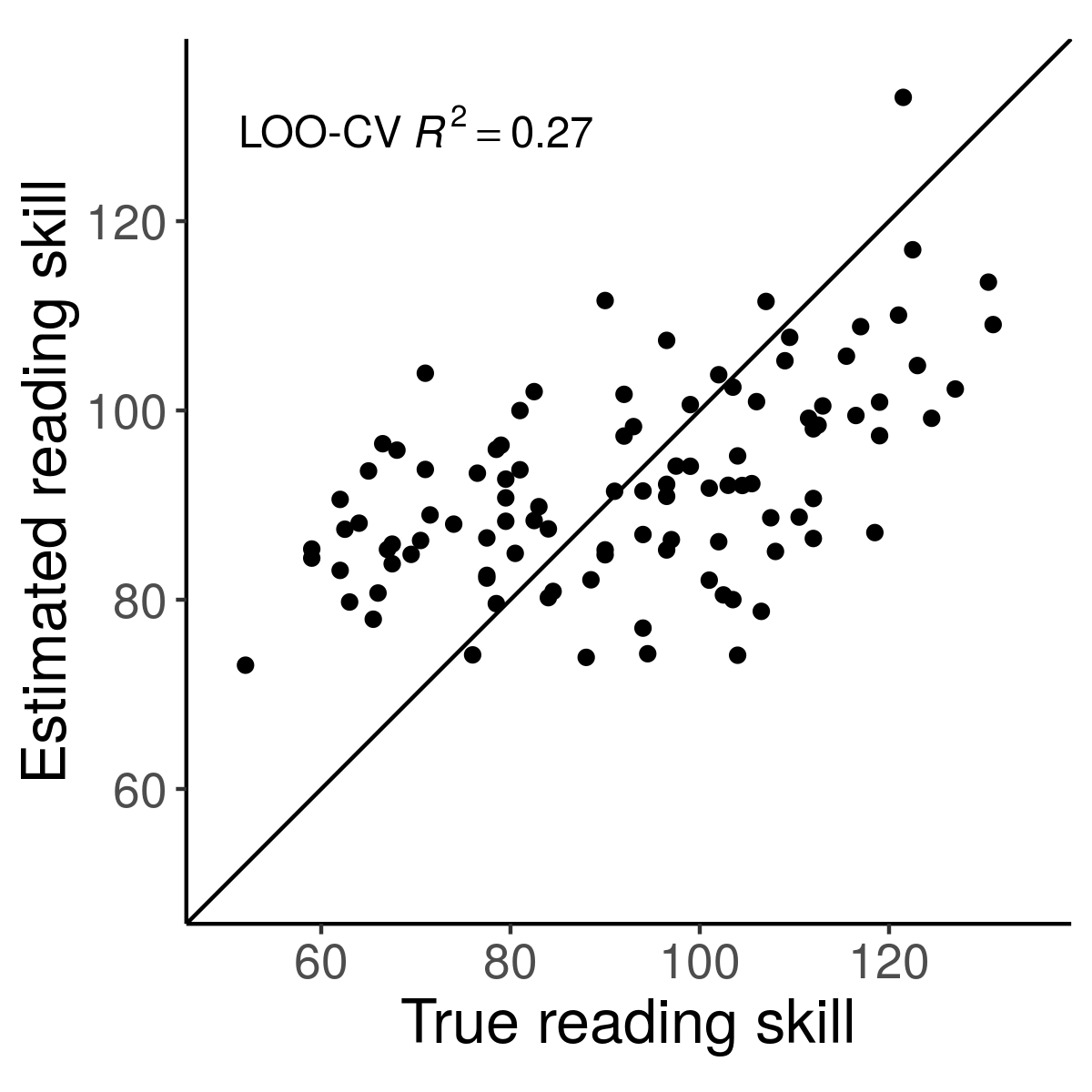

**Figure S6.** *Comparison of true versus predicted reading skill for the single-factor model. Point estimates are computed with leave-one-out cross validation (LOO-CV).*
